# Selected Laboratory Equipment and Kitchen Utensils as an Alternative Sterilization Method for Sterilizing Plant Tissue Culture Media

**DOI:** 10.1101/2023.10.26.564287

**Authors:** Reniel S. Pamplona, Christopher DV Dela Cruz, Maria Angie Tayangona

**Affiliations:** Institute of Crop Science, College of Agriculture and Food Science, University of the Philippines Los Baños

**Keywords:** plant tissue culture, sterilization, low-cost, kitchen utensils, remote learning

## Abstract

Plant Tissue Culture (PTC) is defined as a technique in culturing different plant parts *in vitro* under sterile conditions, and supplemented with optimum nutrients. Due to COVID-19 pandemic which forced everyone to isolate and lock up themselves inside their respective homes, this PTC, which is a heavily laboratory-dependent activity of the students, professors, researchers, and other PTC enthusiasts, was halted. As a response to continuing the PTC remotely, the alternative options for sterilizing PTC media using inexpensive and commercially available laboratory equipment (UV ray and microwave) and kitchen utensils (steamer and casserole) were explored in this study. The commercially available 30mL hinge sauce cup was selected as an *in vitro* media container. The addition of the autoclavable plastic or ziplock significantly reduced the deformity of the containers, and contamination rate of the media. Among the laboratory set-up, the two-minute exposure of PTC media to the microwave produced almost the same sterile media in the autoclave method (control). On the other hand, both boiling, and steaming methods had no significant difference compared with the control. Among all the treatments, the steaming method for 20 min had maintained sterile cultures for 3 weeks, and consistent pH like in the control, making it highly recommended for sterilizing PTC media at remote places, particularly at home. Thus, the alternative sterilization protocol of PTC media has been established through this study.

## INTRODUCTION

The outbreak of the coronavirus or better known as COVID-19, started last December 2019 at Huanan Seafood Wholesale Market, in Wuhan, Hubei, China, where the said illness was associated first to pneumonia because of the symptoms like fever, fatigue, dry cough, and gastrointestinal problems (Huang C, *et al.,* 2020; Wu Y, *et al.,* 2020). Eventually, the virus spread around the globe disrupting normal activities, and taking the lives of many people. All countries immediately closed their borders to prevent the spread of the virus. The citizens of the respective countries were isolated, and mandated to stay at home. In the Philippines, the government halted all outdoor activities, and implemented a quarantine period on March 17, 2020 (IATF, 2020). All establishments except the supermarkets were closed down. This includes schools, universities, and research institutions. Research was frozen for some months, and the education system shifted to remote learning, making it impossible to conduct laboratory experiments that are relying greatly on laboratory set-up.

Plant tissue culture (PTC) is one of the laboratory experiments that need advanced laboratory equipment to execute. This field of science grows explants or plant parts inside a sterile container under an aseptic environment and procedures. The explants are kept inside the media, which are being supplemented by balanced amounts of nutrients and hormones. Aside from the sterilization of the explants, the preparation and sterilization of media through steam, ultrasonic, and heat, is crucial to ensure that no contaminants will grow inside the culture vessel (Misra, and Misra, 2012).

The workplace and the tools should be maintained to utmost cleanliness at all times to avoid the microbial contaminations such as bacteria and fungi. There are some practices that are being implemented to maintain a conducive environment for PTC experiments. Modern laboratories use a special equipment called “laminar flow hood”, where a HEPA filter and a moderate wind pressure reduces the entrance of microorganisms. The application of 95% alcohol, 15 minutes before the activity in the laminar flowhood in the surface of the laboratory tools, equipment, and even to the hands/gloves of the researcher has been proven to prevent the contaminants (Misra, and Misra, 2012). The overnight or limited duration of exposure of the laboratory tools and laminar flow hood to ultraviolet (UV) rays also kills the possible contaminants in the culture. There are also laboratories where the UV light, which is installed in the laminar flow hood, is being widely used for sterilization of laboratory tools and equipment. Although this is harmful when in contact with a person’s skin, the UV light kills the bacteria and spores of fungi that are present in the surface table of the laminar flow hood. In addition to this, the use of UV rays in explant, affecting the physiological growth, and chemical, metabolic and molecular composition of the plants (Cavallaro *et al*., 2022; Craver *et al*., 2014; Klein, 1967; Metwally *et al*., 2019; Pacaldo, and Arradaza, 2021; Tasheva, *et al*., 2015; Vanhaelewyn *et al*., 2020). Thus, this is being discouraged for the sterilization process of explants, and only applied to laboratory equipment and PTC media. The surface sterilization using UV light usually takes 1 hour to 24 hours depending on the laboratory protocol. For laboratory tools such as the forceps, scalpel with blades, scissors, glass bottles, autoclavable plastics, rubber bands, papers, distilled water, etc., It is highly advisable to autoclave these tools prior to the use in the experiments. Autoclaving is an effective method of sterilizing the laboratory equipment to kill the contaminants. The laboratory tools and solid media are generally being sterilized under 121°C for 15 minutes (Howie, 1959; Misra, and Misra, 2012). On the other hand, Burger, D. W. (1988) optimized the protocol for 10 and 4000 ml liquid media under 121°C for 40 min. The sterilization process was being conducted using a pressure cooker only.

In the Philippines, clean empty jars of condiments, sandwich spread, jams, and ketchup are commonly used in PTC laboratories since the material can withstand the high temperature of the sterilization. Aside from the glassware, researchers used materials made of polypyrene, polymethylpentene, polyallomer, Tfzel, ETPE, Teflon FEP (Misra, and Misra, 2012). In addition to this, instead of the metal lid, autoclavable plastic and rubber bands are being used to seal the bottle. This reduces the cost of the *in vitro* container. On the other hand, collecting or buying these materials, particularly bottles/jars, will be challenging for immediate use in the remote set-up. The bottles also require a spacious cabinet or growth chamber for storing the cultures. The need to look for an alternative container that is durable and cost-effective remains as an option for the researchers. In addition to this, there are numerous studies conducted to alter the sterilization process specifically for the PTC media. This includes the use of microwave oven, chemical reagents such as chlorine dioxide (ClO2), Diethylpyrocarbonate, sodium hypochlorite (NaOCl) (Youssef and Amin, 2001; Tisserat *et al.,* 1992; Duan, *et al.,* 2019; Cardoso, 2009; Cardoso and Inthurn. 2018; Macek *et al*., 1995; Gavilan, 2018). Other methods for sterilizing PTC media are not yet explored, and have not been conducted in the Philippines.

In this paper, the use of the easily commercially available laboratory equipment and kitchen utensil are explored to sterilize the plant tissue culture media. As of the researchers’ knowledge, this is the first time that the experiment was conducted in the Philippines. The result will be beneficial to researchers, students, and PTC enthusiasts in doing their research in their respective places.

## MATERIALS AND METHODS

### Observation period and Data Collection

All experiments were conducted at the Plant Tissue Culture Facility. Media under different sterilization conditions were kept at the Plant Growth Chamber with 25°C and 14 hr photoperiod. All data were recorded every week.

### Selection of the Media Container

All experiments were done using commercially available and easy-to-obtain material like 30mL sauce cups, with the dimension of 5cm (top) x 2.5cm (h) x 4cm (bot). The 30ml sauce cup is widely utilized at the fast-food chain, local restaurants, and even in the common home kitchen. The three most common 30mL hinged cups were assessed first before proceeding to the next experiment. Initial materials were obtained from fast food chains, and cleaned thoroughly prior to the experiment. For the main experiments, all containers used for the following experiments were procured online to ensure their sterility and uniformity.

### Sterilization of the MS Media

In order to identify alternative sterilization methodology for remote set-up, the most common kitchen practices plus some easy techniques are listed as different treatments (Table 1). After the selection of *in-vitro* containers, autoclavable plastic or ziplock were used in all of the treatments or sterilization procedures except in Treatment A2 (UV light).

**Table 1.** Treatments for the sterilization of Plant Tissue Culture MS Media. (A) Using common laboratory equipment; and (B) Using Kitchen Utensil.

| Set-up | Number | Treatment | Duration (min) | Brief Description of Procedure |
| --- | --- | --- | --- | --- |
| **A** | 1 | Autoclave (control) | 20 | Pressure cooker or Autoclave; 15 PSI, 120°C for 20 minutes |
|  | 2 | UV light | 10, 20, 30 | Inside the laminar flowhood, closed hinged cups were exposed under UV light. Covered with black cloth for protection. |
|  | 3 | Microwave | 2 | Microwaving of hinged cups under whirlpool microwave oven for 2mins with first run of 1min duration, and in second and third run 30sec durations was done. Moreover, 5min rest interval between the runs were executed to subside heat. |
| **B** | 1 | Autoclave (control) | 20 | Pressure cooker or Autoclave; 15 PSI, 120°C for 20 minutes |
|  | 2 | Steamer | 10, 15, 20 | Steam the samples. Start the timer after the water reach the boiling point. Use food thermometer for monitoring. |
|  | 3 | 70% Alcohol + boiling | 20 | Samples should tightly sealed and stable before submerging to water. Start the timer after the water reach the boiling point. Use food thermometer for monitoring. |

To evaluate the treatments to produce free-from-contaminants media, every 30mL sauce cup was dispensed with 10mL MS Media, and optimized the condition and set-up prior to the main experiment (Figure 1). Generally, the MS media possesses complete nutrients and supplements that can help to maintain an explant or callus *in vitro.* Failure in the sterilization process among the treatments will result in extreme proliferation of pathogens or contaminants in a very short time.

**Figure 1.**
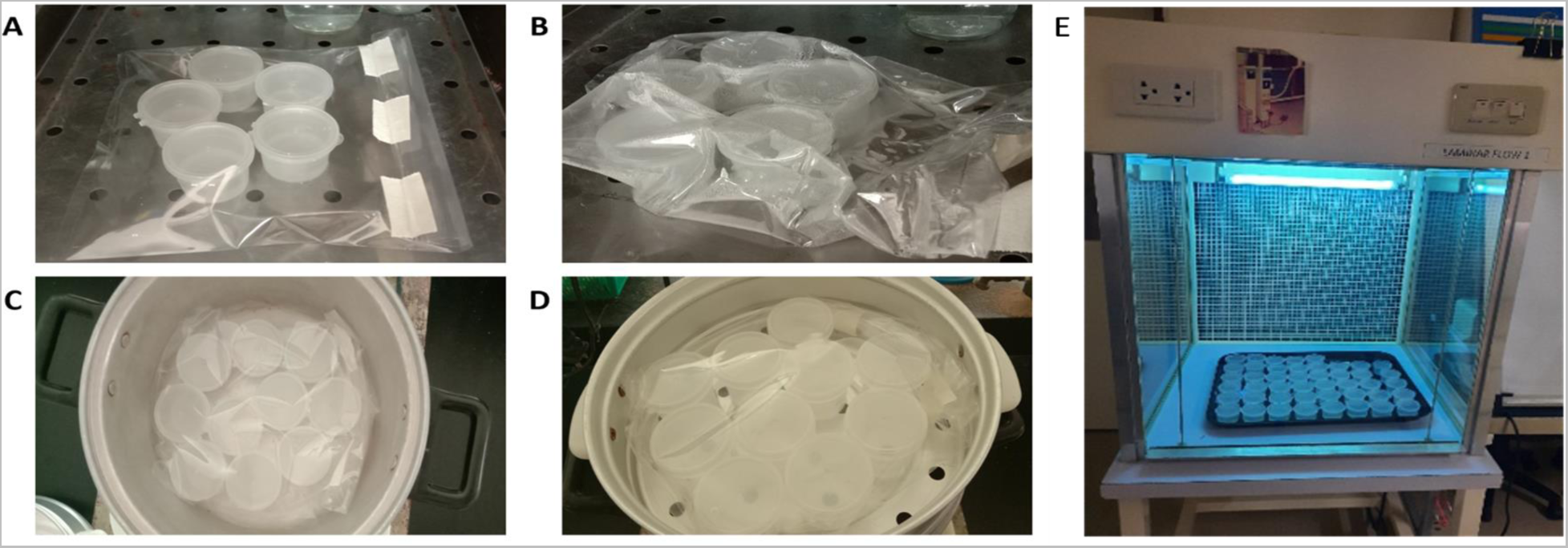
Representative pictures of the experimental set-up for the sterilization process of PTC media. (A) Samples before the optimization experiment; (B) Samples subjected to sterilization using steaming; (C) Samples subjected to 70% Alcohol swabbing of containers + boiling; (D) Samples subjected to steaming; and (E) Exposure of media to UV light.

The autoclave method is used as a standard for this experiment. All treatments had 15 samples with 3 replicates.

### Measurement of pH

To monitor if there were drastic changes in the acidity and basicity of the media, the pH of all treatments were recorded before and after the sterilization procedures using the pH meter. Gelling agents were not included in the media to evaluate only the change in the solution.

### Monitoring of Temperature

To evaluate the effectiveness of the temperature in the sterility of each media, the specific temperature of the treatments were monitored using the food temperature purchased online. Table 2 shows the accurate temperature per treatment and duration.

**Table 2.** Monitored Temperature per treatments using laboratory and kitchen utensils.

| Set-up | Number | Treatment | Duration (min) | Temperature |
| --- | --- | --- | --- | --- |
| A | 1 | Autoclave (control) | 20 | 121 °C |
|  | 2 | UV light | 10, 20, 30 | - |
|  | 3 | Microwave | 2 | 500 – 800 watts<br>800Watt sec = 0.421252<br>Celsius Heat Unit |
| B | 1 | Autoclave (control) | 20 | 121 °C |
|  | 2 | Steamer | 10 | 61.80 ± 0.3464 |
|  |  |  | 15 | 98.83 ± 0.7572 |
|  |  |  | 20 | 100.73 ± 0.2082 |
|  | 3 | 70% Alcohol + boiling | 10 | 101.33 ± 0.5774 |
|  |  |  | 15 | 100.47 ± 0.1528 |
|  |  |  | 20 | 100.47 ± 0.1528 |

### Statistical Analysis

All treatments were conducted in 3 replicates with 15 samples each. Significance of the data collected were evaluated using an R software program using complete random design.

## RESULTS AND DISCUSSION

### Selection of Alternative Media Container

Figure 2 shows the different sauce cups obtained from different restaurants, and subjected to standard protocol for autoclaving. All cups were filled with water and autoclaved with and without a clean autoclavable zip lock. In the preliminary experiment, the sauce cups in Figure 2B were the only samples that didn’t crumple during the sterilization process without a clean autoclavable zip lock. On the other hand, the images depicted in Figure 2A and 2B were placed inside the autoclavable zip lock prior to autoclaving, and showed to withstand the pressure of the autoclave, while the Figure 2C crumpled. The presence of an autoclavable zip lock reduces inside the autoclave machine. For the succeeding experiment, the sauce cup in Figure 2B is selected since it exhibited the durability to endure the high pressure in the autoclave machine. This also conforms to the previous study of Misra, and Misra, (2012) that materials made of polypyrene can be autoclaved and used as containers for PTC. Thus, the material will be able to withstand the other sterilization process in this study.

**Figure 2.**
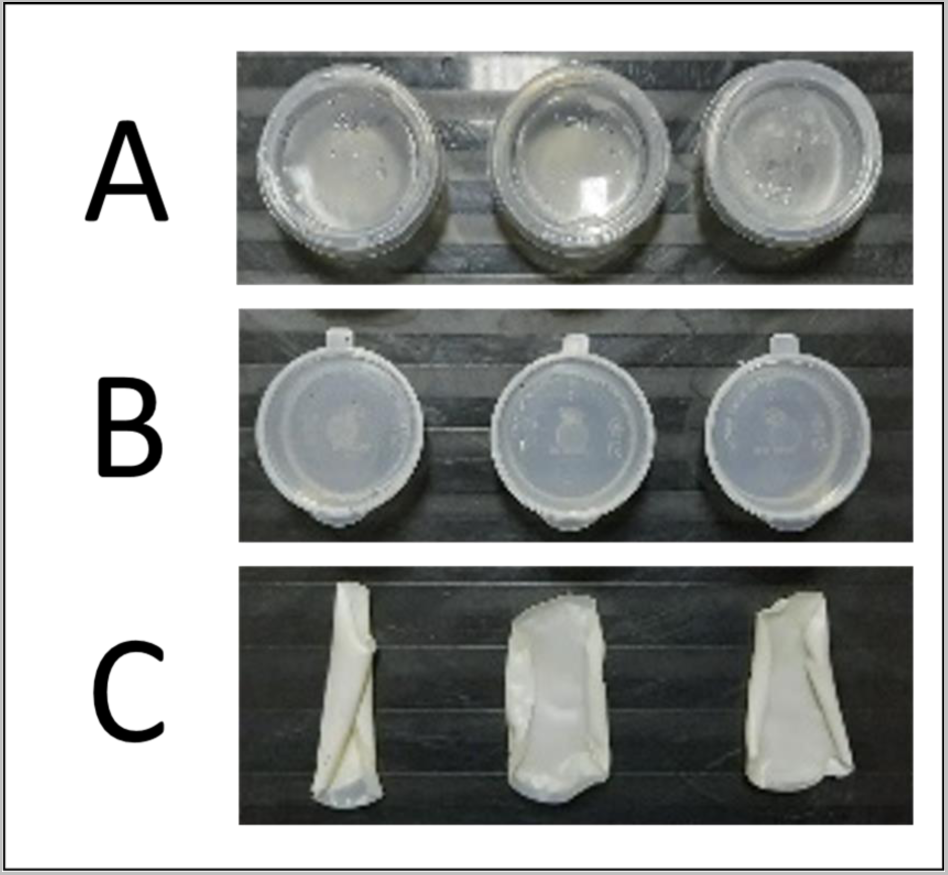
Image of the different 30 mL sauce cups autoclaved inside the autoclavable plastic. (A) Transparent sauce cup with thin structure; (B) Transparent sauce cup with thick and sturdy structure; (C) Opaque plastic sauce cup.

### pH monitoring of the Media

In order to know if there are drastic changes in the acidic or basic property of the nutrients after the sterilization process, the pH level before and after the application of different treatments was also monitored (Figure 3). Most of the PTC protocol maintains the pH of 5.2 to 6.0 prior to autoclaving (Skirvin *etal.,* 1986; Beyl, 2011). But after sterilizing the media using autoclaving, the pH of the media resulted in 5.2 to 5.8 (Figure 3), which is almost similar in the previous study of Msogoya, *et al*. (2008). Aside from the sterilization protocol, the other factors causing changes of pH are attributed to basal medium, carbohydrate source e.g. sucrose, fructose, gelling agent, activated charcoal, and medium storage (Owen, et al., 1991; Msogoya, *et al*. 2008). The changes in the pH can cause vitrification or develop plantlets to have a turgid, glaucous, and watery appearance (Ding, et al., 2008; Shi, et al., 2017). The pH below 4.5 prevents the growth of the plants due to destabilizing of growth regulators, and causing precipitation of phosphate and ion salts (Dodds and Roberts, 1990). This will result in liquefaction of the media. However, Figure 3 also depicts that all of the media under the different treatment produced only solid media. Thus, no significant change happened after sterilization.

**Figure 3.**
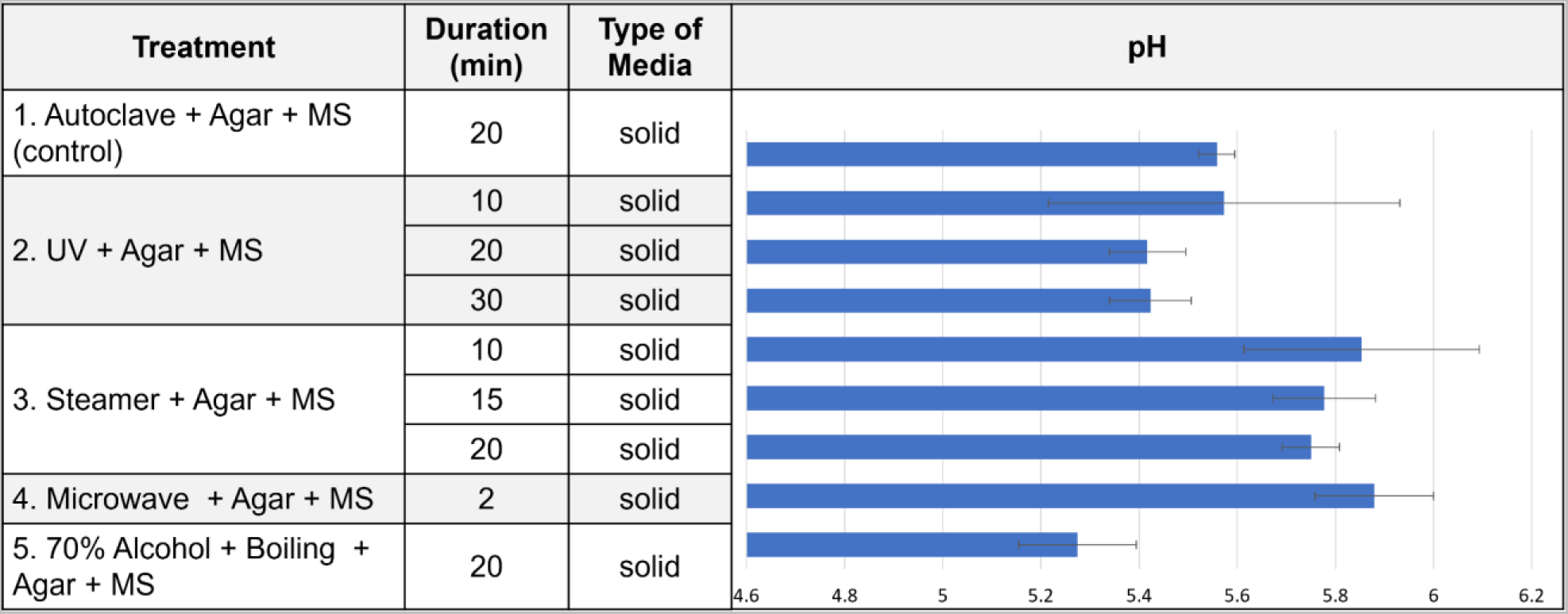
Type of media formed and their corresponding pH Level after sterilization.

### Sterilized Media Containers

The presence of contaminants for 6 weeks were observed in different media containers. Figure 4 shows all the sterilized plant tissue culture media under various sterilizing conditions, while Figure 5 presents the rate of contamination rate per treatment. Both set-ups produced a 100% sterilized media for control (autoclave). In Figure 4.A, media under the microwave sterilization (treatment A3) produced 80% of the contaminant-free media starting week 2 to week 6. According to the previous study of Tisserat et al., (1992), and Youssef, E.M.A. and Amin, G.A. (2001), the microwaved culture media can produce 96%-100% sterilization rates under the condition: 15 min, 390 W or for 5, 10 or 15 min at 520 W. In the experiment, due to the small dimension of the hinged sauce cups and inclusion of autoclavable plastic, the circulation of heat inside can cause an abrupt explosion within 1 min. Thus, the previous study can possibly be achieved in larger containers. Despite the lower sterilization rate, the result still shows a promising possible sterilization process of the media in the remote set-up. On the other hand, the sterilization under UV light (Treatment 4.A2) in different durations produced approximately 0-60% sterilization rate. Thus, this is less effective than the microwave sterilization.

**Figure 4.**
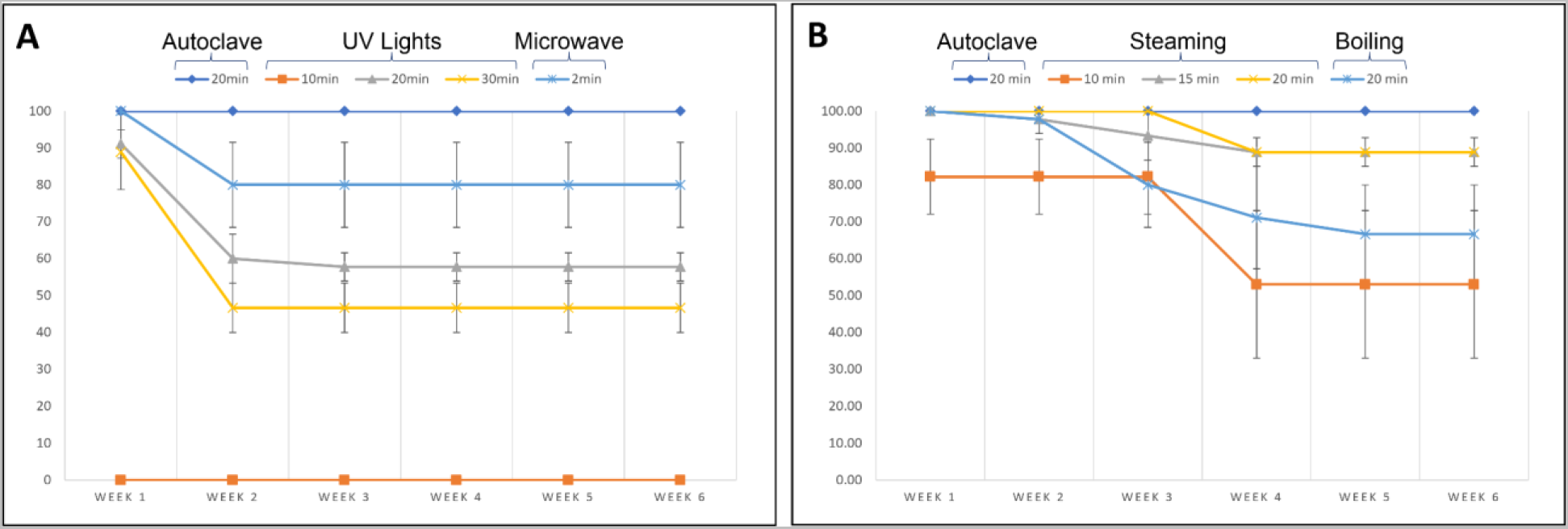
Rate of the Contamination-free Plant Tissue Culture Media after the Alternative Sterilization Procedure. *(A) Using selected Laboratory Equipment; and (B) Using selected Kitchen Utensils*.

**Figure 5.**
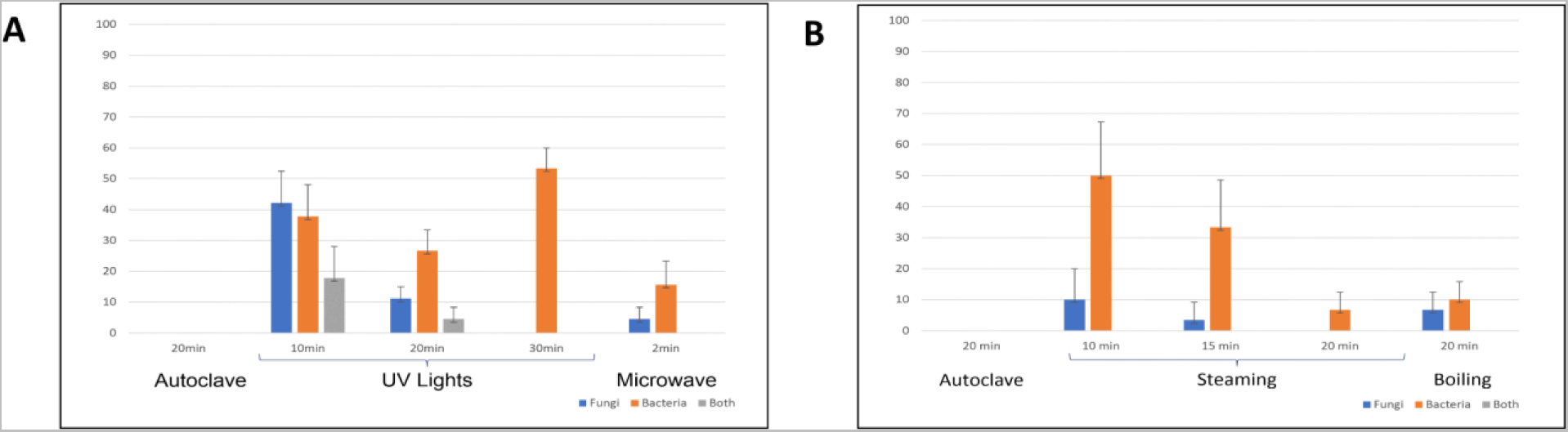
Rate of contamination per treatment after 6 weeks of observation. (A) *Contamination observed from the alternative sterilization protocol using laboratory equipment; and* (B) *Contamination observed from the alternative sterilization protocol using kitchen utensils*.

In Figure 4.B, Both Treatment steaming (B2) and boiling (B3) showed as sterile as the control (Treatment B1) results. Steaming for 15 and 20 min showed the highest sterile media. However, between the two, the treatment B2 for 20 min, maintained a 100% contamination-free until the 3^rd^ week observation. In Figure 5, the most common contaminants in the experiment set-ups were bacteria, which are produced by accidentally transmission of contaminants during the sterilization procedures (Boxus P., and Terzi J.M., 1987; Scortichini, M. and Chiariotti, A. 1987; Wilson C.M., 1966). This was least observed in Treatment B2-20 min, which appeared on the 4^th^ week of the observation. Normally, the media for PTC were changed every 2^nd^ or 3^rd^ week to assure its optimum nutrient content. Thus, treatment B2-20 min or steaming process using the normal steamer is the most highly recommended among all treatments in both set-up A and B as an alternative sterilization method for remote PTC experiments.

## CONCLUSION

In this study, we have established a substitute effective protocol for sterilizing plant tissue culture media. Readily available food containers made of polypyrene were validated to serve as good containers for *in vitro* culture. The smallest container such as a 30 mL sauce cup can be used for remote PTC. Additionally, the use of autoclavable plastic or ziploc can reduce the deformation of the containers, without compromising the sterility of the media. Among the treatments, the steaming for 20 min can be an alternative way for sterilizing PTC media. The media are free from contaminants until week 3. Mostly the contaminants are bacteria which can be attributed to the aseptic techniques. Results showed that the treatments didn’t affect the solidification, and pH-level. The study can be adapted for remote learning or in research in their respective residence.

## CONTRIBUTIONS

RSPamplona conceptualized and designed the experiments. RSPamplona and MADTayangona performed the experiment and did the data collection. RSPamplona and CDVDela Cruz validated the research design, performed the data analysis, and wrote/revised the manuscript. All authors proofread and approved the manuscript.

## ACKNOWLEDGMENT

The authors would like to extend its deepest gratitude to UPLB-OVCRE Basic research fund. We would also like to thank Mr. Mark Anthony T. Ramos, and Ms. Meldy Reyes for helping in setting up the experiment. Moreover, we are also extremely grateful to Ms. Regina S. Caunin for her administrative support.

